# Multidisciplinary approach to the study of artworks: environmental conditions, color palette, microbial colonization and nannofossils in *Las Musas* painting

**DOI:** 10.1101/2022.10.05.511059

**Authors:** P. Calderón-Mesén, D. Jaikel-Víquez, M.D. Barrantes-Madrigal, J. Sánchez-Solís, J. Mena-Vega, J. Arguedas-Molina, K. Ureña-Alvarado, G. Maynard-Hernández, L. Santamaría-Montero, M. Cob-Delgado, E. Angulo-Pardo, Felipe Vallejo, M. Sandoval-Gutiérez, A. M. Durán-Quesada, M. Redondo-Solano, O.A. Herrera-Sancho

**Affiliations:** Centro de Investigación en Estructuras Microscópicas, Universidad de Costa Rica, 2060 San Pedro, San José, Costa Rica; Facultad de Microbiología, Universidad de Costa Rica, 2060 San Pedro, San José, Costa Rica; Centro de Investigación en Enfermedades Tropicales (CIET), Universidad de Costa Rica, 2060 San Pedro, San José, Costa Rica; Escuela de Química, Universidad de Costa Rica, 2060 San Pedro, San José, Costa Rica; Instituto de Investigaciones en Arte, Universidad de Costa Rica, 2060 San Pedro, San José, Costa Rica; Escuela de Ingeniería Eléctrica, Universidad de Costa Rica, 2060 San Pedro, San José, Costa Rica; Escuela de Física, Universidad de Costa Rica, 2060 San Pedro, San José, Costa Rica; Escuela de Artes Plásticas, Universidad de Costa Rica, 2060 San Pedro, San José, Costa Rica; Department of History of Art, Cornell University, Ithaca, NY, 14853; Instituto Costarricense de Investigación y Enseñanza, en Nutrición y Salud, 42250 Cartago, Costa Rica; Instituto de Investigaciones en Estratigrafía (IIES), Grupo de Investigaciones en Estratigrafía, y Vulcanología (GIEV-Cumanday) y Departamento de Ciencias Geológicas de la Universidad de Caldas, Calle 65 # 26-10, 1700004 Manizales, Colombia; Departamento de Geología, Facultad de Ciencias, Universidad de Salamanca, España. Plaza de los Caídos, s/n, 37008 Salamanca; Escuela Centroamericana de Geología, Universidad de Costa Rica, 2060 San Pedro, San José, Costa Rica; Departamento de Física Atmosférica, Oceánica y Planetaria, & Laboratorio para la Observación del Sistema Climático, Escuela de Física, Universidad de Costa Rica, 2060 San Pedro, San José, Costa Rica; Centro de Investigación en Ciencias Atómicas Nucleares y Moleculares, Universidad de Costa Rica, 2060 San Pedro, San José, Costa Rica

**Keywords:** analytical methods, dyes/pigments, tangible, cultural heritage, oil paintings, art conservation

## Abstract

Over time, cultural heritage has become a key for comprehending and developing our society at an individual and group level, as it provides fundamental information about our origins, specific temporary space, materials’ availability using current technology, artist’s intention, and site weather conditions. Here, we investigated the physical characteristics of an Italian large-format artwork diptych, located on the ceiling of the National Theater of Costa Rica, to evaluate its pictorial palette as well as the tropical climatological conditions and the fungal aerial spore concentration. We characterized the role of two innovative software tools, as they have direct connection with an effective microbiological sampling and description of secrets encompassed in each pictorial layer during the process of creation and intention. We further found that eight genera of calcareous nannofossils could be associated with the optical properties of the artwork and the effects that the artist wanted to portray through his creative process.

## Introduction

Since ancient times, humanity has produced works of art and faced various difficulties related to the production, conservation, and restoration of what we now call cultural heritage. The ultimate quest to reconstruct the object, its history, the artist’s intention, its color changes or structural integrity, and the impact due to climate changes and historical moment lies not only in having a realistic understanding of the dynamics of the object’s materials but also the storage conditions, habitat conditions, along with the interaction between the object and an entire environment during a time frame. For this reason, the history of pictorial materials and their artistic techniques involves experimentation and creativity in the artistic and scientific fields, as these areas and their characteristics should be considered while exploring the object^[1]^. The paintings, known as *Musas I* and *Musas II*, decorate the ceiling of the space originally used as Men’s Canteen room at the National Theater of Costa Rica (NTCR) (built in 1891-1897). They embody the Greek god Apollo and the Muses at Mount Parnassus for our artistic visualization) ^[2]^. The two canvases were painted in Milan in 1897 by Carlo Ferrario (1833-1907), an Italian painter, theater decorator, and teacher who worked at the Academia di Brera as well as at the Teatro alla Scala^[3]^. The methodological results obtained from the scientific analysis conducted in these paintings have been contrasted with historical data about the materials used by Carlo Ferrario ^[4,5]^. Valuable information was found in regard to the improvement of public policies for the conservation and restoration of artworks in latitudinal position.

The artworks are unique along with their history, the artist’s intention, the dynamics of their materials, and their interaction with their surroundings. However, large-format paintings have limited access, and this affects the acquisition and comprehension of information in a non-invasive manner^[6,7]^. To conduct a nonintrusive approach, a technical examination of the artwork and its environment is essential. Moreover, application of multi-analytical studies -as the ones previously performed in the paintings of the NTCR^[8–12]^-; extensive monitoring of temperature, humidity, and atmospheric pollutants (as fungal spores and carbon dioxide, CO_2_) in the space where the artworks remain; isolation and identification of microorganisms (fungi and bacteria) that inhabit the surface of the paintings were the methods executed in order to determine if the physical-chemical properties of the artworks were affected by microorganisms and climate conditions^[13–15]^.

Normally, paintings consist of many superimposed layers where the pigments and the binder interact causing color changes and discoloration. These are inherent to the artist’s selection of materials and the restoration interventions. Moreover, this multiplex structure changes linearly over time. Some of these changes can be attributed to microorganisms and weather conditions (spores, temperature, humidity, and CO_2_), as they can accelerate those processes ^[16,17]^. We employed imaging techniques in combination with a state-of-the-art software in order to elucidate the apparent color of the artworks based on distribution, arrangement, and size of each pigment. The aim of this approach was to determine the average separation distance between ultramarine blue pigments because our previous research on the color palette used by the artist revealed that this one was the predominant pigment ^[8]^. On the other hand, strong evidence of microbial contamination was found on *Musas I* and *Musas II*. For this identification, we applied a software that uses UVF photography to distinguish possible areas of microbial growth in the paintings. The reliability of its measurements has an accuracy of approximately 11%, corresponding to a high-efficiency quantitative determination. This assessment provides an exciting opportunity to advance in our interpretation of an early detection of microbial contamination, while using a software as well as a systematic and minimally invasive approach.

As the materials in oil paints not only interact with each other but also with their surroundings over time, we need to analyze the artworks similar to a dynamic process. *Musas I* and *Musas II* consist of oil paint over hemp canvas with approximately 125 years of fabrication ^[9]^. They have four well defined layers: top paint layer, intermediate paint layer, second ground layer, and first ground layer ^[8]^. These components have been gradually changing due to their interaction with biotic and abiotic factors, as they affect the conservation of the painting and the perception of it.

Ground layers, also known as *gesso*, are preparatory and priming steps for the later oil-bound layers ^[18]^; they are a combination of calcium carbonate materials, glue, and white pigments. In the case of *Musas I* and *Musas II*, the white pigments used by the artist were zinc white and lead white^[8]^. The calcium carbonate materials, known as chalk, consist of deep marine deposits, and they are characterized by the presence of calcareous nannofossils^[18,19]^. The study of the biodiversity and abundance of these nannofossils could provide information about the geographical distribution of these organisms^[20,21]^, and therefore, it can provide clues to identify the origin of the materials. Ground materials and white pigments could have different optical properties ^[19,22]^, and the fabrication process of lead white could result in subtypes of the crystalline lead carbonate in the pigment. Some of the forms might be cerrusite and hydrocerussite. They change the opacity and brightness of the painting ^[22]^. The ground layers in combination with the colored pigments layers and the canvas materials create a complex stratigraphic structure that composes the artwork. In general, therefore, these findings regarding ground layers enhance our understanding of the nature of the artworks.

This study delves into the analysis of the materials (type and distribution) used in the paintings *Musas I* and *Musas II*, the environmental conditions surrounding them, the interactions of microorganisms with the artworks, and the paintings’ state of conservancy. The study’s objective was to examine an oil painting on canvas diptych from an unique interdisciplinary perspective, as it implements history, art, and applied sciences to cultural heritage in order to analyze the composition of substances that the artist employed to create the paintings and the journey of the artwork - considering that it was created in Italy and later transported to Costa Rica- to provide tools for the study of priority areas of intervention to conserve this cultural heritage.

## Results and Discussion

### In search of the origins of the artworks

Carlo Ferrario, a well-known theater decorator, painted the two canvases with pigments that were available in the Milan art market during the late nineteenth century, as they came from various European mines and paint manufacturers ^[1–4]^ (Figure 1). We have verified that this diptych contains lead red, viridian, ultramarine, vermilion, chrome yellow, zinc white, and lead white^[8]^. Most of these pigments are recommended by Ferrario in his treatise *La tecnica della pittura ad olio ed a pastello*, in which he reveals the working methodology and formulas that he prefers in order to produce a wide variety of colors in oil painting ^[5]^. According to a bibliographic and geographical review, the source material for the main pigments came from different possible mines around the world (Figure 1B, the location of the mines is indicated in figure caption). The lead and zinc used in the lead white and zinc white as well as other pigments have a close relation with galena minerals and their derivatives ^[23]^. Moreover, they can came from other mines in Italy such as Distrito Minero SW Sardinia^[24,25]^, Distrito Minero SE in Sardinia^[26,27]^, Salafossa^[28,29]^, and Rabil Cueva del Predil^[30]^. Sources for red pigments mainly came from mercury, as they are associated with the Cinabrio mineral. This mineral probably was extracted from two mines in Italy: Mt. Amiata and Abbadia San Salvatore ^[31,32]^. Nonetheless, other mines could have been involved in its extraction such as Almadén, Spain^[33,34]^ and Idrija, Slovenia^[35,36]^. In the case of the blue pigments, they were obtained from the Lazurite mineral. Moreover, the most probable mines for this mineral are Sar e Sang, Afghanistan ^[37]^; Baikel Lake Slyudyanskii, Russia^[38–40]^; and Flor de los Andes, Chile^[41]^. Finally, to clarify, Ferrario used synthetic ultramarine pigments instead of natural ones^[8]^.

**Figure 1.**
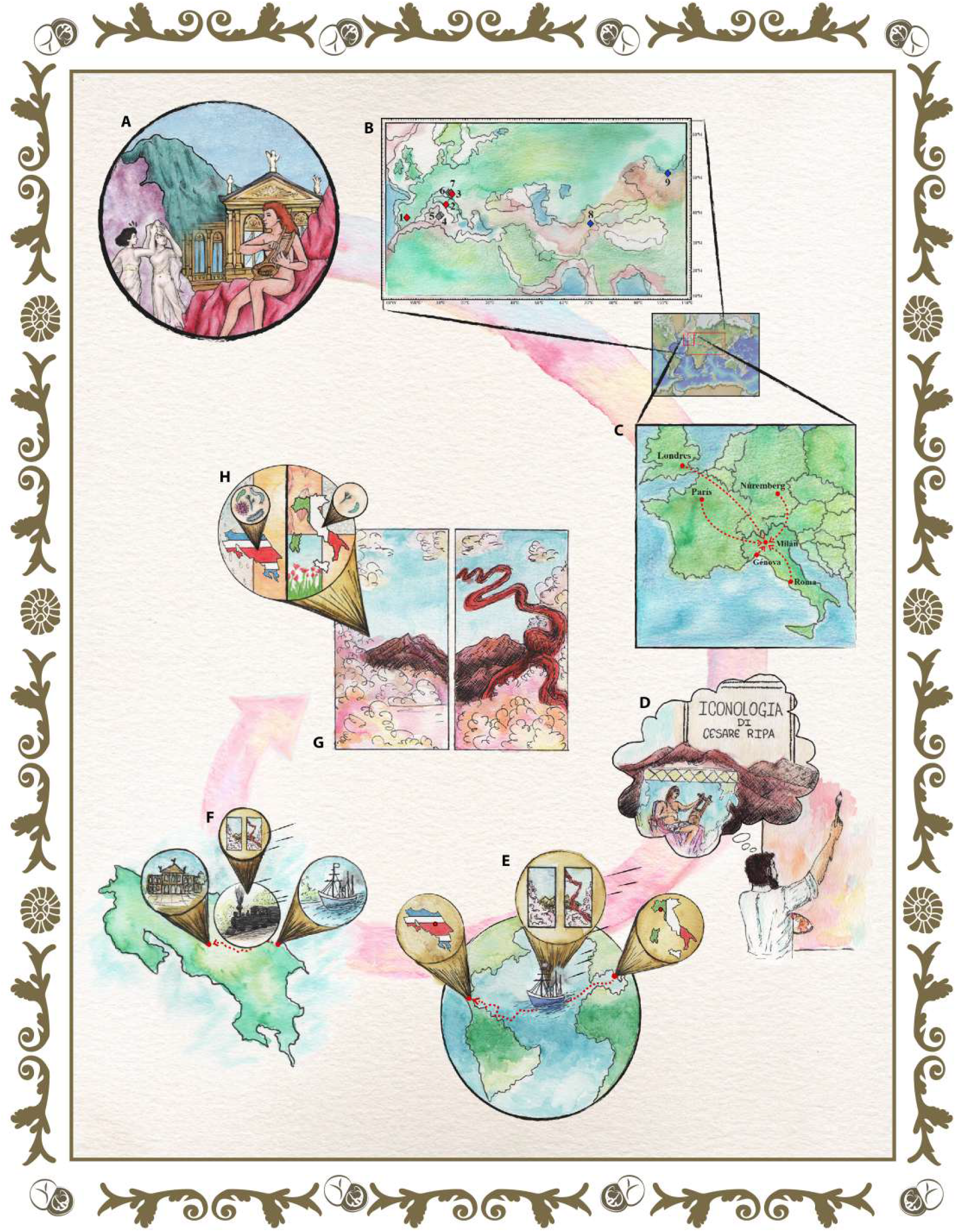
From the origin to the present of paintings. To understand the context of the *Musas I* and *Musas II* oil paintings (**A**), this study includes information of (**B**) the probable mines from which the minerals were extracted for the pigments’ elaboration used in the paintings, (**C**) the trade routes that took them to Milan, where the artist Carlo Ferrario created the artworks, inspired by classical iconographic representations such as those established by Cesare Ripa (**D**). Subsequently, we discover that the works were transported to Costa Rica by sea (**E**) from the port of Genoa to Puerto Limón, (**F**) where they were subsequently transferred by rail to the National Theater of Costa Rica, in the country’s capital, San José. There those paintings (**G**) still decorate the Theater even though weather conditions differ from those in Europe. This difference in the environment surrounding the paintings has led to their present deterioration, also produced by the presence of contrasting microorganisms (**H**).

*Musas I* and *Musas II* were commissioned by the Government of Costa Rica in 1896 and, they were painted by Ferrario in Milan in 1897^[2]^. The iconography of Ferrario’s diptych materializes the legacy of Cesare Ripa’s Iconologia (1593) and the influence of Neoclassicism in the representation of Apollo and the Muses at Mount Parnassus^[2]^. Historically, this iconographic theme has been considered suitable for an opera house due to the association of the muses with artistic inspiration. Moreover, in ancient Greek culture, Apollo is the god of the arts; his aesthetic influence is evident in the NTCR (Figure 1D). Ferrario’s paintings were shipped from Italy to Puerto Limón, Costa Rica. Then, a few months before the NTCR inauguration in 1897, they were transported to San José by railway (Figure 1, E and F). However, Ferrario’s paintings brought to Costa Rica not only classical iconographies and European pigments but also nannofossils older than Mount Parnassus itself (see below).

### Current location and state of conservation

#### Environmental conditions

The assessment of current damage and risk of damage of cultural and artistic assets related to climate conditions is often considered from the perspective of works that are directly exposed to the environment, such as historic buildings and statues; nonetheless, this topic is becoming relevant within the context of temperature and humidity changes that is associated with the climate change^[42]^. The impact of environmental changes is also present on the inside of buildings, threatening artworks and driving attention to consider the influence of climate variability and change in conservation^[43]^. In tropical areas, where rainfall variability and warm temperatures favor moist environments, the risk of damage of cultural and artistic heritage is even higher. Moist environments in buildings with limited isolation enable the propagation of spores and provide suitable conditions for microbial development. The NTCR is in San José province -an area that exhibits an annual rainfall exceeding 1800 mm/year on average and a mean temperature ranging between 14 °C and 24 °C approximately- (see Figure 2, E, D and C). An abundant rainfall during the year and a marked interannual variability contribute to moist conditions in the vicinity of the NTCR. Changes in the space not only are limited to climate variations but also to a specific anthropogenic pressure associated with increasing combustion of fossil fuels. The growth in the number of vehicles that circulate the roads surrounding the NTCR contributes to a rise in the pollution inside the building. Moreover, these meteorological conditions favor the deposition of atmospheric pollutants. Hence, to deliver adequate solutions for the conservation of the artworks that are part of the NTCR, it is of great importance to monitor the external and internal environmental conditions in order to have a better understanding of how these factors are compromise them.

**Figure 2.**
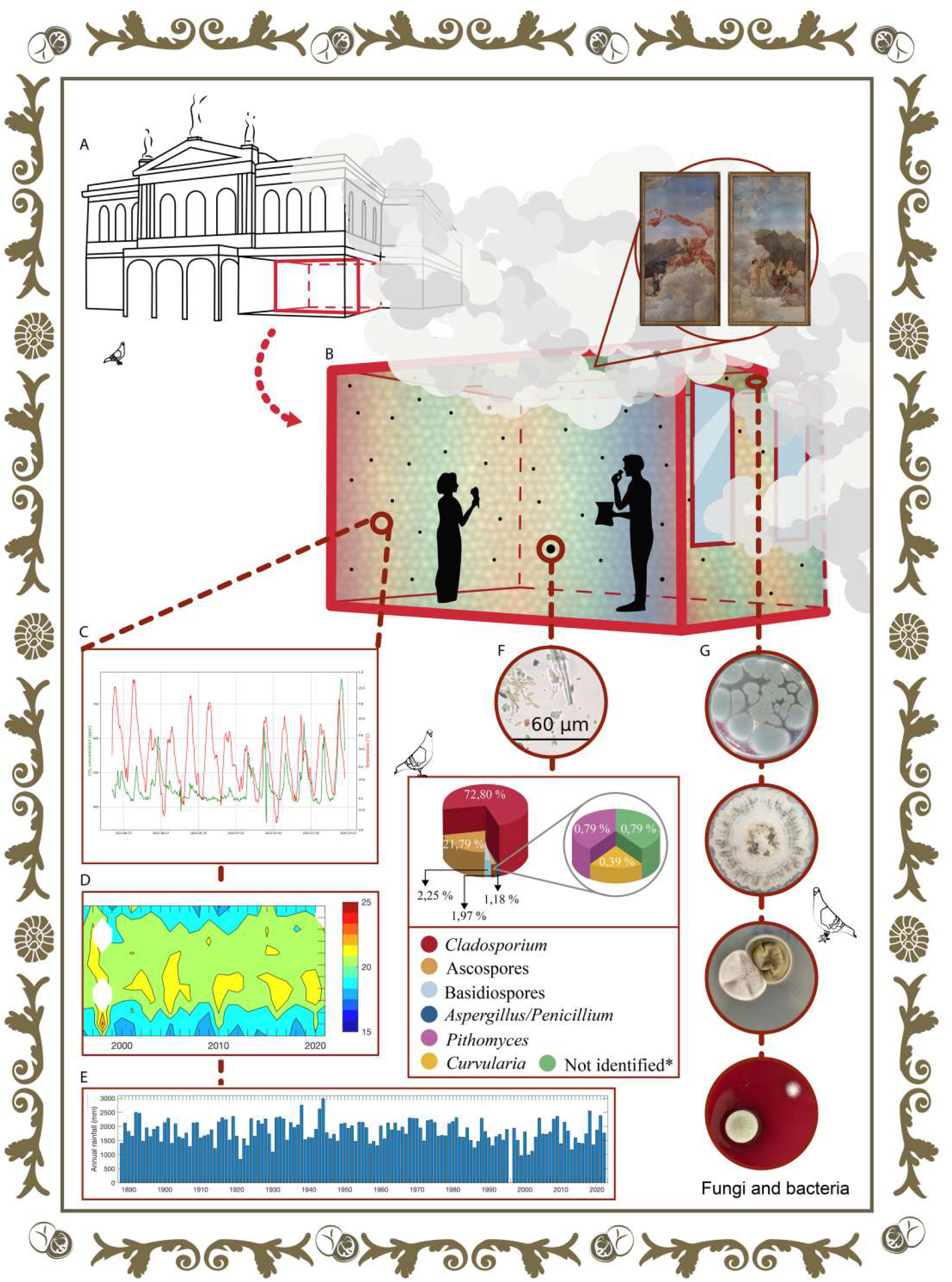
The tropical environmental conditions of the pictorial diptych living day by day. (**A**) The NTCR building is located in the capital of the country, in a very busy place characterized by a constant flow of vehicles and people. (**B**) *Las Musas* reside in the ceiling of the former Men’s Canteen. (**C**) shows the monitoring of the environmental conditions (temperature, humidity, and CO2) for a period of three months. The temperature variation since the year 2000 is shown in (**D**). The amount of precipitation per year is observed in (**E**). A fungal aerial volumetric study was conducted in this room, where *Cladosporium* was the main fungal spore present (**F**). Finally, (**G**) shows some previously isolated and characterized bacteria and fungi from the paintings (from top to bottom: *Penicillium* section *Chrysogena*, Hyaline fungi, *Cladosporium halotolerans* with *Cladosporium dominicanum* and *Cladosporium halotolerans* with *Penicillium crustosum*). The regions of microbiological interest were previously selected by means of the computational tool *MicroorganismPattern* and UVF photography.

Figure 2 reveals a steady and constant shape of the air temperature throughout the time that our stations monitored the environmental conditions. The monitor was specially designed for the examination of this diptych. The diurnal cycles of this variable reach their maximum around 15:00 hour and a minimum around 3:00 hour. For this *in situ* study, measurements were collected for approximately two months. This study aims to appreciate if the demeanor of the variable is typical or atypical. The NTCR was compared with other museums such as the Louvre Museum, in Paris; the Metropolitan Museum of Art, New York; and National Palace Museum, Taipei, Republic of China. This comparison showed that the highest mean air temperature corresponds to the NTCR (23.5 °C) followed by the National Palace Museum (21 °C)^[44]^, Louvre Museum (12 °C)^[45]^, and finally, the Metropolitan Museum of Art (12 °C)^[46]^. In the case of the relative humidity variable, we found almost the same sequence of results: National Palace Museum (82%)^[44]^, followed by NTCR (72.7%), Metropolitan Museum of Art (67%)^[46]^, and Louvre Museum (65%)^[45]^. The most surprising result that emerged from the analyzed data was regarding the carbon dioxide (CO_2_) concentration variable. The National Theater of Costa Rica has the highest value for this variable: 570 ppm. The order for the other places is as follows: Metropolitan Museum of Art (465 ppm), National Palace Museum (423 ppm), and the Louvre Museum (414 ppm). The high value for CO_2_ can be attributed to the vehicles that transit the roads surrounding the NTCR as well as to people that visit the room where the paintings reside. These findings need to be cautiously interpreted, as the sample size was a clear limitation. Finally, there is abundant room for future research in this area.

#### Microbiological detection with computer program

The large-format diptych (total area of roughly 36.5 m^2^) is an object that is alive and, at the same time, hard-to-reach for art professionals and restoration scientists. The large size and location (on the ceiling of the Men’s Canteen) of the artworks complicate the search for microorganisms. The most successful restoration and conservation approach is the one that follows a non-invasive technique such as systematically selecting regions of interest for microbiological sampling by means of software tools. For this reason, with the help of a novel software (*MicroorganismPattern*) along with UVF images, we identified places with possible bacteria and fungi colonization. The software uses an algorithm that is based on a template matching method^[47]^. First, we analyze UVF photographs (Figure 3B) in order to find a characteristic perceivably pattern and locate it. This pattern that resembles a probable type of deterioration, is established through cautious observation of the UVF photography of the artwork. The UVF image and the microorganism pattern are exemplified as matrix *I* and matrix *R* in panel C, respectively.

**Figure 3.**
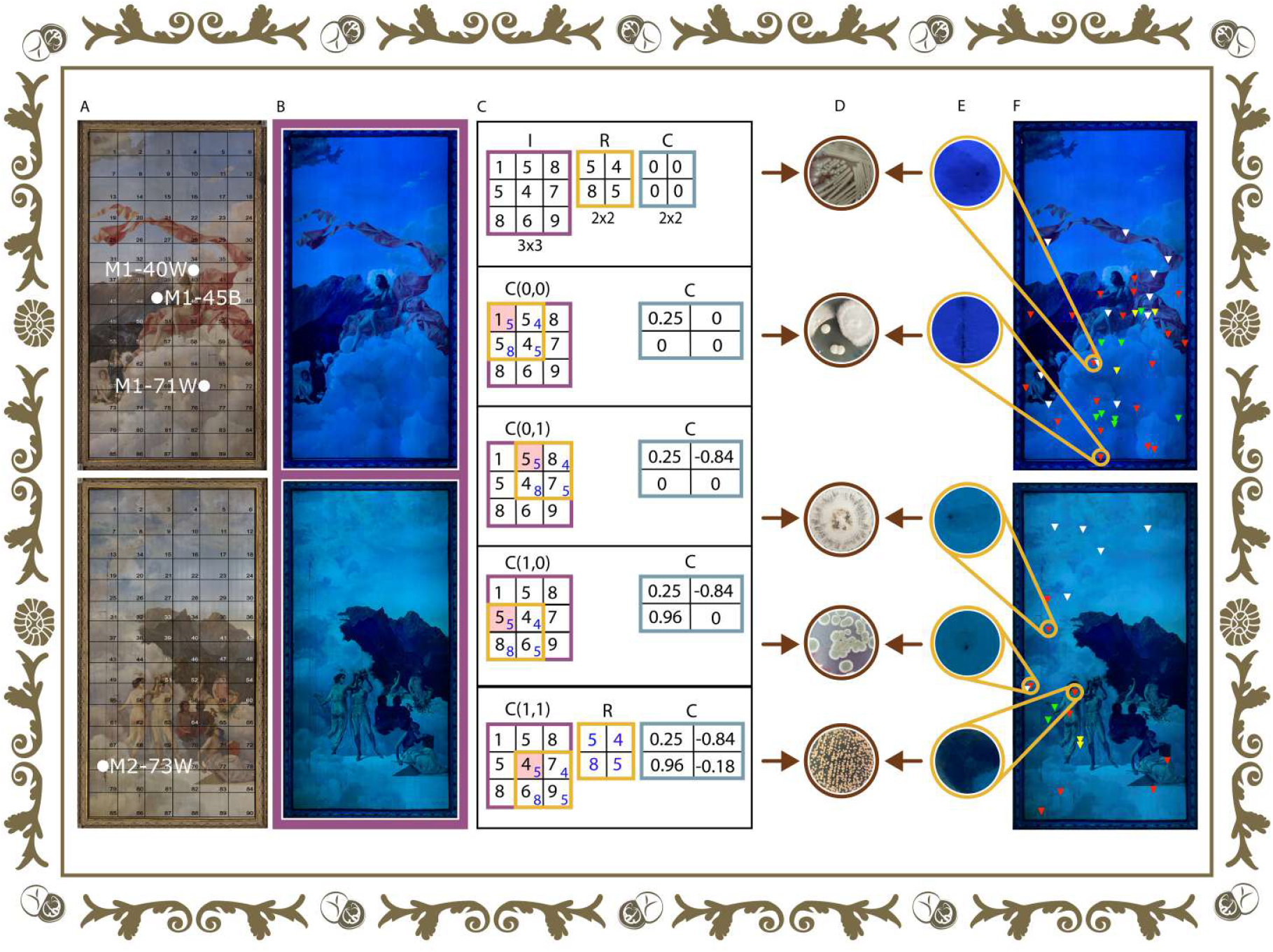
Determination of the biodeterioration areas of interest for direct microbiological sampling. (**A**) Photographs of the paintings *Musas I* (top panel) and *Musas II* (bottom panel) in visual light with information of the samples of painting layer extracted for microscopical analysis.(**B**) Photographs of the paintings *Musas I* and *Musas II* in ultraviolet fluorescence (UVF) light. (**C**) First, the UVF photograph shown in (B) is discretized by its pixels and the information placed in the matrix called *I* (see purple color). As described in the text, via meticulous observation, possible deterioration patterns are selected and placed on the artwork (see E). These patterns are used as a basis in the template matching method to carry out the object detection, and the pixels of the *R* matrix are accordingly assigned (see yellow color). Finally, the *MicroorganismPattern* computational tool estimates matrix *C* using Equation 1, as explained in Methods. From the example indicated here, the value of the matrix *C* that is close to the value +1 is the input *C*(1,0). (**D**) Microorganisms isolated from the deterioration areas (from top to bottom: *Bacillus subtillis*, Fungi from family Polyporaceae, Hyaline fungus, *Penicillium steckii, Cryptococcus uniguttulatus*) **(E)** Examples of significant types of deterioration observed in the painting (from top to bottom: types of deterioration); the areas were determined with UVF images. (**F**) Location of biodeterioration areas according to software (white) and *Musas I* visual biodeterioration areas: rounded dark blue (red), rounded light colored (green), and rounded fluorescent (yellow); as for *Musas II*, visual biodeterioration areas: rounded faint colored (red), rounded light colored (green), and rounded dark blue (yellow).

Next, a matrix *C* (Figure 3C) is computed by the method of correlation coefficient between the pattern (R) and the entire artwork (I) (see Methods for details). As a pilot study, we established the regions of interest through the accessible multidisciplinary software *MicroorganismPattern*. Then, we carried out two microbiological samplings in the large-format diptych. The most significant finding of this systematic sampling was that the software had a high-efficiency determination of approximately 11%; however, it is important to bear in mind that all the localities discovered were not sampled. From thirty-one areas of around 1 cm^2^ that were swabbed, eighteen microorganisms were isolated and identified (see below).

#### Microbiological findings

In general, the paintings from the NTCR are exposed to constant interaction with human and environmental factors like relative humidity, temperature, and light that induce the deterioration processes ^[16]^. Therefore, it is expected that a significant number of microorganisms may colonize and, eventually, deteriorate the general structure of each artwork^[48]^. The study of the role of some microbial populations that are present in these paintings is imperative for further elaboration of proper conservation strategies.

In both *Musas I* and *Musas II*, specific physical patterns that may correlate with deterioration were identified (see Figure 3 for the location of the sections). A total of nineteen samples –thirteen during the first sampling date (including the negative control) and six in the second– were collected from *Musas I*. In the case of *Musas II*, thirteen samples (twelve during the first sampling date and one from the second) were gathered. From the first sampling date, three bacteria (*Alicyclobacillus acidoterrestris, Bacillus cereus*, and *Kocuria rizhophila*) and one fungus (*Penicillium* section *Chrysogena*) were isolated from *Musas I*; the fungus and two of the bacteria were obtained from the same sampling area (87D3). On the other hand, from the second sampling date, one bacterium (*Sphingomonas paucimobilis*), seven filamentous fungi (two *Cladosporium halotolerans*, two basidiomycetes from family Polyporaceae, one *Cladosporium dominucanum*, one *Penicillium crustosum*, and a the yeast *Rhodotorula glutinis*) were isolated.

Moving forward to *Musas II*, three fungi (*Penicillium steckii*, *Cryptococcus uniguttulatus*, and a non-identified hyaline fungus) and one bacterium (*Bacillus cereus*) were obtained from the first sampling period. In the case of the second sampling date, only one bacterium (*Staphylococcus lugdunensis*) was isolated. No microbial contamination was detected on the negative control site. It is imperative to mention that different sampling areas were selected for each period.

*Penicillium* was the main fungal genus present in the paintings, as it was found in both sampling periods and both artworks, followed in importance by *Cladosporium* (found just in one sampling date). Both genera are widely described as contaminants of closed indoor environments ^[48]^. The results obtained are coherent with the aerial volumetric examination at the Men’s Canteen (location of the paintings). In this room, the main circulating fungal spores were conidia from the genus *Cladosporium* (1293 spores/m^3^), *Ascospores* (387 spores/m^3^), Basidiospores (40 spores/m^3^), and conidia of either *Aspergillus* or *Penicillium* (21 spores/m^3^) (cf. supplementary material). Also, it is worth noting that these fungi have enzymes (cellulases) ^[49,50]^ that might be able to hydrolyze the components on the canvas and the wood from the paintings’ frame. Interestingly, one yeast was present in *Musas II (C. uniguttulatus*). This fungus is a normal inhabitant of pigeon’s (*Columba livia*) droppings^[51]^. Hence, its presence may be related to the park areas that are close to the NTCR. The blastospores may have been transported by wind to the paintings.

On the other hand, the isolation of bacteria confirms that the process of microbial decay in the artwork may be a complex one, as it may involve the interaction of different microbial communities; the findings from the 87D3 spot supports this idea (this sample area is the one in the botton border of *Musas I*, see Figure 3F). All the bacterial genus that were identified (*Bacillus spp., Alicyclobacillus spp., Kocuria spp., Staphylococcus spp*., and *Sphingomonas spp*.) are microorganisms commonly present in the environment, as they have capacity to form endospores (*Bacillus spp*. and *Alicyclobacillus spp*.) ^[52]^ or biofilms (*Kocuria spp., Staphylococcus spp*., and *Sphingomonas spp*.) ^[53]^. Given that the surface of the canvas and artwork may offer limited supply of nutrients and humidity for bacterial growth and survival, it is not surprising that bacteria with the ability to form resistant phenotypes are the ones obtained from the paintings.

Interestingly, *S. paucimobilis*, a bacterium commonly reported in clinical settings, was isolated from *Musas I*. This bacterium is characterized by a strong adhesion capacity to synthetic surfaces due to its ability to produce exopolymers^[54]^. Other members of the *Sphingomonas* group have been found in ancient wall paintings ^[55]^, but to our knowledge, this is the first time it has been reported from canvas. Similarly, *K. rizhophila* is increasingly recognized as an emergent human pathogen with high capacity to colonize abiotic surfaces ^[56]^. Moreover, this species has been related with biosorption of heavy metals from soils, suggesting that it may adapt to extreme environments ^[57]^. *Kocuria* and related species have also been previously isolated from art pieces ^[58]^.

In the case of *B. cereus* and *A. acidoterrestris*, they are spore forming Gram positive bacteria with capacity to produce extracellular degradative enzymes. *B. cereus* has been widely known as a foodborne pathogenic and a spoilage microorganism able to synthesize proteases, lipases, lecitinases^[59]^, quitinases ^[60]^, and pectinases ^[61]^. Similarly, *A. acidoterrestris* is recognized for producing decarboxylases that spoil acidic fruit juices ^[62]^. Nonetheless, this is the first report on *A. acidoterrestris* colonizing an artwork. Both microorganisms could be able to survive in the limiting conditions offered by the paintings’ surface, and they may eventually cause some type of damage to them. In fact, *B. cereus* has been previously isolated from XVII century paintings. It is speculated that it may be responsible for decoloration and other types of alterations ^[63]^. However, additional experiments are necessary to elucidate if these bacteria and other microorganisms are able to have a significant impact on the conservation process of the patrimonial paintings from the NTCR.

#### Microscopic study of the structure of the artworks and return to origins

Aside from the history and environmental and biological conditions, the comprehension of the paintings’ materials and structure is therefore a crucial aspect that completes our insight of these artworks. Our previous studies provided us with information about the layers and their composition ^[8]^; nonetheless, we carried out a further investigation with the aim of finding more secrets of these paintings.

Regarding the painting layers of *Musas I* and *Musas II*, in this work, we sought to establish a proof-of-concept methodology for the understanding of the relationship between the observed color and the distribution of the pigments in the color layers. As can be noted in Figure 4, the proposed procedure consists of the observation of the cross-section of a specific sample (M1-45B) directly extracted from the painting (Figure 4A). In the case of M1-45B, it has a dark blue coloration to the naked eye, as it appertains to the paintings’ “mountain area”. For the process, we created a 3D representation of the samples previously studied. This graphic source allows us to outline the information related to the macroscopic color and the main materials found in that particular area of the painting ^[64]^, see Figure 4A.

**Figure 4.**
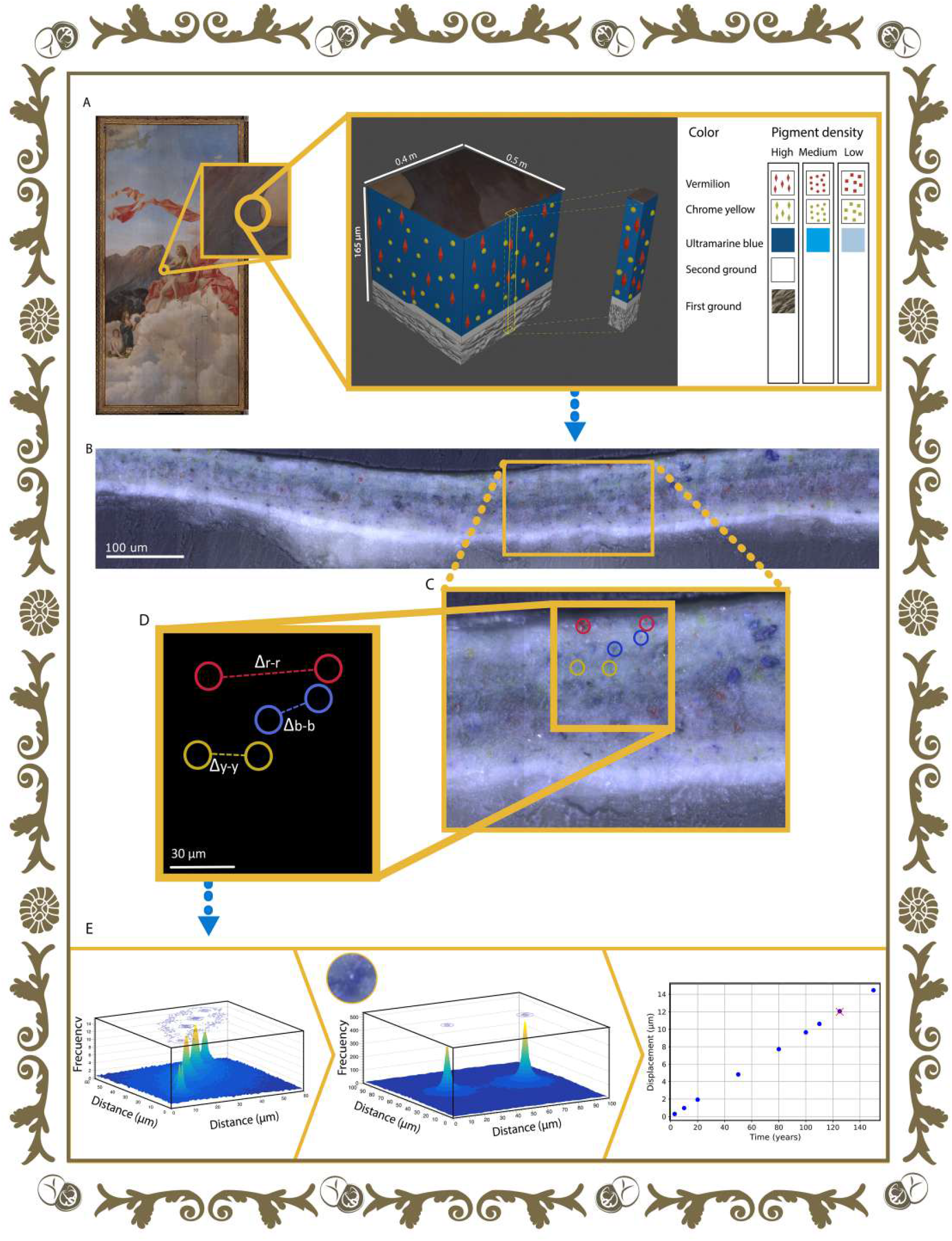
Scheme of the study of the distance between pigments crystals. **A**) First, a sample is extracted from *Musas I* aiming to observe it with the microscope in order to understand the stratigraphy of the painting, as represented in a 3-coordinate system. (**B**) Then, the microscope images of a region of interest of the cross-section of the sample are used as input into the software. (**C**) The software recognizes crystals by color and carries out (**D**) measurements of the distance between crystals of the same color (Δb-b for blue crystals, Δr-r for red crystals and Δy-y crystals), and finally, providing (**E**) histograms of the results along with the theoretical analysis of blue pigments per year.

The upper layer corresponds to the paint layer. In this case, the layer is composed of ultramarine, chrome yellow, and vermilion crystals. Below this layer, it is found the second ground that principally contains lead white. The latter pigment has different compositions (hydrocerussite and cerussite), and each one generates distinct effects in the painting. This has been observed in famous works such as Vermeer’s *Girl with a Pearl Earring* ^[22,65]^. Finally, the first ground layer consists of chalk; later, we will detail this layer (Figure 4B).

Moving forward, the color layers are a mixture of binder and pigments that provides color to the artworks. The software developed, called *PigmentsSeparation*, analyzes a selection of photographs that were acquired by microscope (Figure 4C). *PigmentsSeparation* differentiates the different colors of crystals (e.g., blue, red, and yellow) and measures the distance between them (Figure 4D and E). As a result, histograms of the distances between crystals of the same color (e.g., blue-blue) and different colors (e.g., blue-red) are obtained (Figure 4E).

The average distance between the blue crystals, red crystals, and yellow ones can be observed in the histograms in Figure 4E. It was expected that blue crystals were going to have the shortest distances between themselves. The reason behind this is that the blue crystals are the ones with a higher density. The result was an average distance of 29 *μ*m (standard deviation of 12 *μ*m) between blue crystals. This matches the hypothesis because the distance for blue crystals is shorter than the distance between the yellow and red crystals, as they have 31 μm (standard deviation of 16 *μ*m) and 41 **μ**m (standard deviation of 13 **μ**m), respectively. As expected, the color present in a larger quantity and with shorter distance between themselves predominates in regard to the color observed macroscopically.

The comparison of the two summaries of the statistics about blue crystals (see Figure 4F) reveals that there is probably crystals dynamics within the layers resultant of gravitational attraction. Interestingly, due to the oldness of these artworks (circa 125 years old), it is possible to assume that the theoretical rheology of the pigments in the binder (linseed oil) can approximate a sphere that falls in a viscous reservoir towards the center of the Earth, as it travels an estimated distance of 12 *μ*m in that same period. In general, therefore, it seems that considering the dimensions of this paint stratigraphy, the drag force given by Stokes’ law^[66]^, seeing the medium as a Bingham fluid (i.e. the viscosity of 1×10^9^ Pa·s) ^[67]^, subsequently, the surprising correlation displacement of the ultramarine pigment can be found in Figure 4E. The findings corroborate the association between the movement of the pigments within the stratigraphy and the date of the painting. The test demonstrates that this proof-of-concept methodology could be used to explore and determine the artist’s intentions and creative process as well as the artwork’s life span by virtue of the pigments motion.

An important limitation of *PigmentsSeparation* is its reduced capacity to recognize pigment crystals. If the paint layer has dark color, difficulties occur when identifying crystals of lighter colors such as yellow. An interrogative arose while analyzing the distances between crystals: have the crystals had this distribution since the painting’s creation? Considering deterioration processes such as the generation of metal soaps in lead pigments, it is unlikely to have the migration and accumulation of material. These two processes are known for changing the microscopic structure^[22,68,69]^ and the optical effect of the painting ^[16,70]^. Our new tool *PigmentsSeparation* might lead studies specialized in monitoring paintings over time. The changes in crystal distribution could be recorded through a systematic treatment. This software is easy to implement for specific solutions. Ultimately, our research is characterized by taking a novel look at the first ground layer, as it applies a new approach for analyzing the origins of the materials used in *Musas I* and *Musas II*.

#### Materials reveals nannofossils and properties in the painting

Calcareous nannofossils are important algal primary producers in marine environments. Their tests (skeletons) have accumulated over the ocean floors since the Jurassic to recent times^[71]^. Chalks are calcareous rocks principally composed of the accumulation of these algae. Moreover, chalks are well preserved and exposed in northwestern European seas. This soft white calcareous rock was commonly used as a primer in historical paintings, as it gave a matte and delicate finish ^[72]^.

The calcareous primer in *Musas I* and *Musas II* was studied, through the analysis of the samples M1-71W, M1-40W, and M2-73W (see sample location in Figure 3) with the Scanning Electron Microscope (SEM), in order to examine the composition of the first layer. The images showed abundant and well-preserved nannofossils of calcareous nannofossils. The nannofossils were classified using the taxonomic guidelines of previous studies ^[73,74]^ and the biostratigraphic ranges of identified taxa, following the standard low-latitude zonation of Sissingh ^[75]^ and Burnett *et al*. ^[76]^. The micropaleontological analysis suggests an age of Late Cretaceous. The calcareous nannofossils assemblage observed in the analyzed images consists of *Cribrosphaerella spp*., *Eiffellithus* spp., *Kamptnerius magnificus, Micula* spp., *Prediscosphaera cretacea, Retecapsa* cf. *surirella, Watznaueria* spp., and *Zeugrhabdotus* spp. (Figure 5). The occurrence of the genus *Micula*, characterized by its first appearance in Biozones UC10 and CC14 and its extinction in Biozones UC20 and CC26, enables us to restrict the age of the material to a biostratigraphic interval ranging from Coniacian to Maastricthian (Figure 5) ^[75,76]^. The identified taxa have a wide paleogeographic distribution. Moreover, they have been reported globally at low, middle, and high latitudes ^[20,77]^. This precludes a more detailed analysis of the paleoecological affinities of the nannofossils, at it could work as an indicator of the origin of the materials used in the paintings.

**Figure 5.**
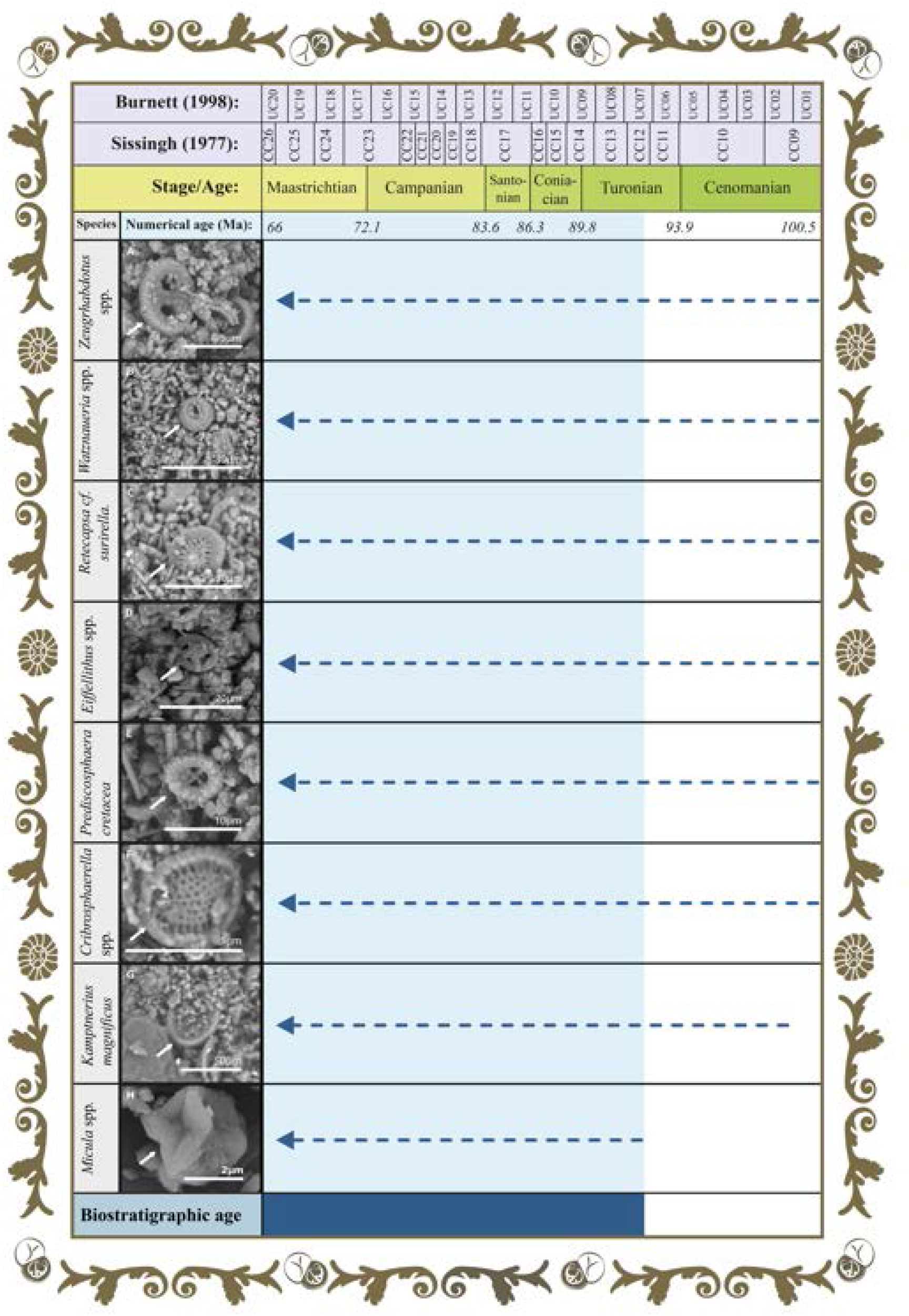
The first ground through the perspective of time: biozonation of the nannofossils of chalk. Biostratigraphic ranges of calcareous nannofossils identified in the analyzed images from *Musas I* and *Musas II* artworks. The species of calcareous nannofossils correspond to: **(A)** *Zeugrhabdotus* spp., **(B)** *Watznaueria* spp., (**C)** *Retecapsa* cf. *surirella*, **(D)** *Eiffellithus* spp., **(E)** *Prediscosphaera cretacea*, **(F)** *Cribrosphaerella* spp., **(G)** *Kamptnerius magnificus*, and **(H)** *Micula* spp. Light blue bar indicates biostratigraphic age.

The samples studied came from areas of the painting where the artist used mainly white pigments (Figure 4A). Calcium carbonate and zinc mainly compose the first ground layer. In the case of the second ground layer, it is primarily constituted of lead white^[8]^. Although the mix of ground material and white pigment makes possible the white tone, it also provides the painting with some optical properties. The bright effect is a result of the chalk being used as a ground material. It is not only related to the amount of coccoliths that constitutes this material but also to the combination of filters and binders^[19,22]^. In the diptych, the predominant color is white^[8]^ possibly because of the artist’s combination of ground with white pigments (lead white and zinc white) and binders. Therefore, we recommend further studies to ascertain the optical effect present in these paintings.

## Conclusion

The eight different species of nannofossils discovered in this study could lead to source of the chalk, as it is of great importance for understanding the properties of the material as well as artist’s intended or desired effect on the painting. In conclusion, an interdisciplinary approach based on composition, novel computational tools (*MicroorganismPattern* and *PigmentsSeparation*), contamination and/or colonization of microorganisms, and state of conservation have been presented in this research, with the aim of exploring the diptych and obtaining new information about the artist’s artistic process. The proof-of-concept methodology proposed for the monitoring of the environmental conditions and fungal aerial spores in the artworks in company with the software *MicroorganismPattern* establishes a high-efficiency and non-invasive approach that could be applied to artworks of worldwide renown. Ultimately, from the experimental data, we try to reveal the detailed nature of the complex artist’s color palette; our evidence provides emerging insight into Carlo Ferrario’s intentions and creative process, as the ground might be the possible combination of itself with eight nannofossils taxa, white pigments, and binder.

## Experimental

Please refer to Supporting Information.

## Acknowledgements

We would like to thank Fabiola Salas Barahona, Plastic Arts School, University of Costa Rica, for her contribution to the artistic design for Figure 1. We give special thanks to Daniel Monge Badilla and Jorge Abarca González, Central American School of Geology, University of Costa Rica, for their valuable support in the material’s geological information. We wish to thank Natalia Rivera Echadi, Bianca Rivera Mejía, and Natalia Camacho Cambronero, Microbiology Faculty, University of Costa Rica, for their support in microbiological analysis. We thankfully acknowledge the support received from Natalia Cordero Villalobos, Sofía Vindas Solano, Meysell Loaiza Brenes, and Karina Salguero Moya, National Theater of Costa Rica, to make this research possible. Finally, we are grateful for the support given by the Vicerrectoría de Investigación, Universidad de Costa Rica to accomplish this research work.

## Entry for the Table of Contents

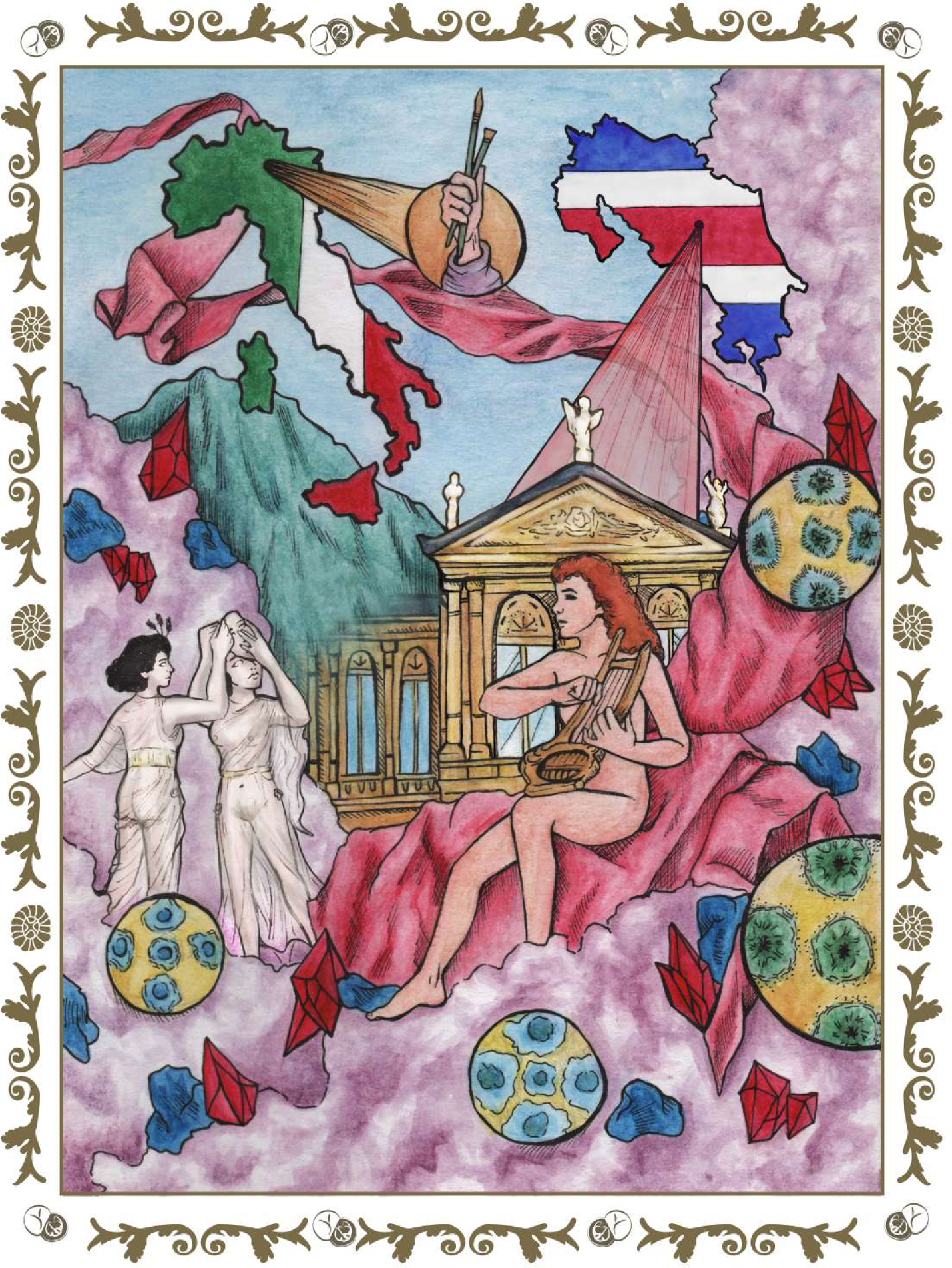
Uncovering the glittering past to cherish and protect the present of cultural heritage. A deeper study of the color palette, environmental conditions, microorganisms as well as nannofossils present in the first ground of the paintings was conducted along with different softwares. This research describes the historical context, origins, and tropicality of a large-format Italian diptych, *Musas I* and *Musas II*, located in Costa Rica’s National Theater.

